# Titin-based force regulates cardiac myofilament structures mediating length-dependent activation

**DOI:** 10.1101/2023.11.09.566413

**Authors:** Anthony L. Hessel, Michel N. Kuehn, Nichlas M. Engels, Devin L. Nissen, Johanna K. Freundt, Weikang Ma, Thomas C. Irving, Wolfgang A Linke

**Author notes:** **Corresponding authors** Anthony L. Hessel,; Wolfgang A. Linke,. Address for both: Institute of Physiology II, Robert-Koch-Str. 27b, D-48149 Münster.

## Abstract

The Frank-Starling law states that the heart’s stroke volume increases with greater preload due to increased venous return, allowing the heart to adapt to varying circulatory demands. Molecularly, increasing preload increases sarcomere length (SL), which alters sarcomere structures that are correlated to increased calcium sensitivity upon activation. The titin protein, spanning the half-sarcomere, acts as a spring in the I-band, applying a SL-dependent force suggested to pull against and alter myofilaments in a way that supports the Frank-Starling effect. To evaluate this, we employed the titin cleavage (TC) model, where a tobacco-etch virus protease recognition site is inserted into distal I-band titin and allows for rapid, specific cleavage of titin in an otherwise-healthy sarcomere. Here, we evaluated the atomic-level structures of amyopathic cardiac myofilaments following 50% titin cleavage under passive stretch conditions using small-angle X-ray diffraction, which measures these structures under near-physiological (functional) conditions. We report that titin-based forces in permeabilized papillary muscle regulate both thick and thin myofilament structures clearly supporting titin’s role in the Frank-Starling mechanism.

## Main Text

The famous Frank-Starling law states that the heart’s stroke volume increases with greater preload due to increased venous return, allowing the heart to adapt to varying circulatory demands. Molecularly, increasing preload increases sarcomere length (SL), which alters sarcomere structures that are correlated to increased calcium sensitivity upon activation. The titin protein, spanning the half-sarcomere (Fig. 1A), acts as a spring in the I-band, applying a SL-dependent force suggested to pull against and alter myofilaments, critically supporting the Frank-Starling effect.^1^ Altered titin-based forces play a crucial role in the etiology of many cardiomyopathies; however, the disease state obscures titin’s role, impeding therapeutic solutions. We solved this problem using the titin cleavage (TC) model, where a tobacco-etch virus protease (TEV_P_) recognition site is inserted into distal I-band titin and allows for rapid, specific cleavage of titin in an otherwise-healthy sarcomere.^2^ Titin cleavage decreases myocardial passive force and stiffness while altering cardiomyocyte tensegrity.^2^ Here, we evaluated the atomic-level structures of amyopathic cardiac myofilaments following 50% titin cleavage under passive stretch conditions using small-angle X-ray diffraction, which measures these structures under near-physiological (functional) conditions.^1^

**Figure 1.**
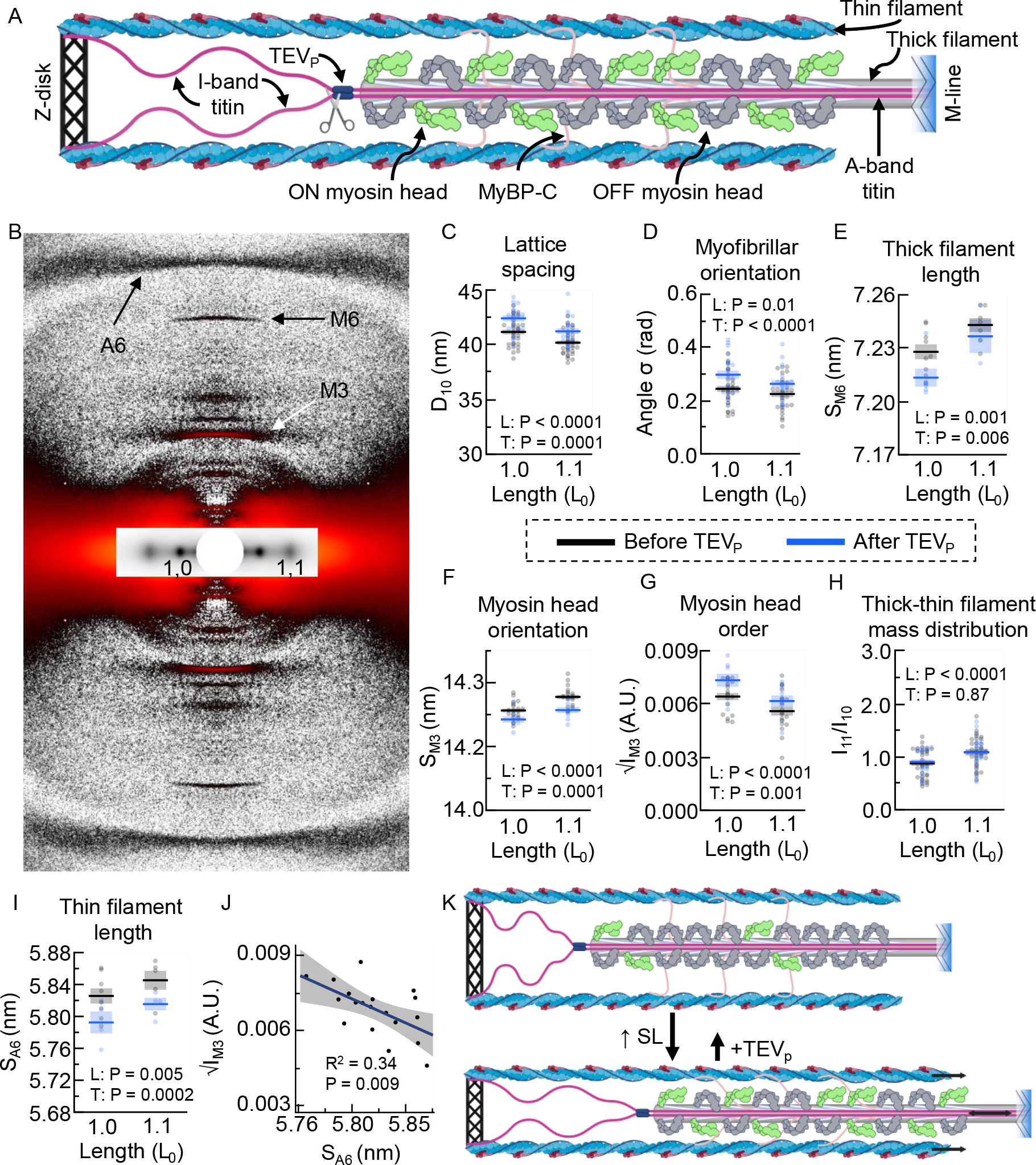
Molecular changes to cardiac sarcomeres after 50% titin cleavage. **A**. Schematic of a half-sarcomere with myosin heads shown in ON (green) and off (gray) states. In the titin-cleavage model, I-band titin is cleavable at the TEV protease recognition site (scissors). **B**. An X-ray diffraction pattern of permeabilized TC papillary muscle, with markers of interest labeled. **C-I**. Analysis of myofilament structures at different sarcomere lengths (expressed as length relative to slack (L_o_)), before (gray) and after (blue) 50% titin cleavage. Both the technical diffraction nomenclature, as well as what they represent, are included. Overlaid are the main effects ANOVA results for length (L) and treatment (T). Interaction main effects were never significant (P>0.05). **J**. Regression analysis between √M3 intensity (√I_M3_) and A6 spacing (S_A6_), with R^2^ and P-value included. **K**. A summary of our findings for sarcomeric structures at short (top) and long (bottom) lengths. We directly demonstrate a relationship between titin-based forces and the myosin head OFF-to-ON transition. Datasets are generated from papillary muscle preparations of 28 heterozygote TC hearts, age range 4-9 months, and presented as mean±s.e.m.

We prepared permeabilized papillary muscle from genotypically heterozygote TC mice (left ventricles) and placed them in a bath of physiological relaxing solution attached to a mechanics apparatus, as reported.^3^ Diffraction patterns (Fig. 1B) were collected from each preparation at an initial length just above slack (100% L_0_; ∼1.9 μm SL)^2^ and at 110% L_0_. The sample was then incubated with TEV_P_ (100 units acTEV in 300 ml relaxing solution) for 20 minutes, sufficient to cleave 50% titins,^2^ rinsed, and the protocol repeated. Diffraction patterns were analyzed using the freeware MuscleX.^1,3^ For each parameter, we used a mixed-model ANOVA, with fixed effects length, treatment (pre/post), interaction, and a repeated-measures random effect (individual). Model assumptions were assessed via residual analysis, the significance level was P=0.05, and all data was reported as mean±s.e.m. For brevity, the main effect P-values are within Fig. 1.

The lateral myofilament lattice spacing is influenced by titin-based forces and was evaluated via the 1,0 reflection, produced from the geometric planes within the thick-thin filament overlap.^1^ Before and after TEV_P_ treatment, the spacing of the 1,0 (d_10_; Fig. 1C) reflection decreased with increasing SL, though the lattice was expanded after titin cleavage, as was expected when some but not all titins are cleaved.^3^ We quantified myofibrillar and myofilament orientation, a determinate of cardiomyopathies,^1^ using the angular spread of the 1,0 reflections (angle σ; Fig. 1D), and found greater disorientation after titin cleavage, highlighting a role for titin-based force in papillary-wide order^.4^

Increasing titin-based forces at longer SLs could stretch the cardiac thick filament, accompanied by a transition of some myosin heads from an OFF to an ON confirmation. This mechanism may increase the myosin head’s ability to form crossbridges, contributing to the Frank-Starling effect.^1^ Here, we show that the spacing of the M6 (S_M6_; Fig. 1E), a measure of thick filament length, increases at the longer SL and decreases with 50% titin cleavage. We further observed myosin head OFF-to-ON transition with increasing SL before cleavage, as indicated by increasing M3 spacing (S_M3_; Fig. 1F; axial periodicity of crowns), decreasing √M3 intensity (√I_M3_; Fig. 1G; proportional to number of ordered heads), and increasing ratio between the intensities of 1,1 and 1,0 reflections (I_11_/I_10_; Fig. 1H; mass distribution between thick and thin filaments). Importantly, 50% titin cleavage decreased S_M3_ and increased √I_M3_, suggesting myosin head ON-to-OFF transitions; however, I_11_/I_10_ presented no change. Therefore, the radial position of myosin heads (I_11_/I_10_) may not change, while myosin heads still reorient (S_M3_; √I_M3_). As a cautionary note, I_11_/I_10_ can be affected by lattice changes, as caused by titin cleavage (Fig. 1C-D). We conclude that reducing titin-based forces leads to a myosin head ON-to-OFF transition while leaving the length-dependent OFF-to-ON transition mechanism intact.

Interestingly, thin filament length, as measured by the A6 reflection spacing (S_A6_; Fig. 1I), also increased with increasing SL, and of note, decreased after titin cleavage, similar to TC skeletal muscle.^3^ The length change of the thin filament is perplexing, as titin is not in an ideal position to stretch it. We postulate that low-level crossbridges in passive cardiac muscle^5^ are recruited from the ON-state motors and transmit strain to the thin filament; less ON-state heads leads to fewer crossbridges and thus shorter thin filaments. This hypothesis is supported by a significant negative correlation between √I_M3_ and S_A6_ (Fig. 1J). The physiological meaning of this phenomenon remains to be elucidated.

In summary, titin-based forces in permeabilized papillary muscle regulate both thick and thin myofilament structures (Fig. 1K), clearly supporting titin’s role in the Frank-Starling mechanism. Further details could be provided using intact TC cardiac preparations, but the experiment requires shuttling TEV_P_ into intact preparations, which is a future goal. A study of TC papillary muscle during contraction is the next logical step.

## Acknowledgments

This research used resources of the Advanced Photon Source, a U.S. Department of Energy (DOE) Office of Science User Facility operated for the DOE Office of Science by Argonne National Laboratory under Contract No. DE - AC02 - 06CH11357, and further NIH support. The content is solely the responsibility of the authors and does not necessarily reflect the official views of the National Institute of General Medical Sciences or the National Institutes of Health.

## Source of funding

German Research Foundation grant 454867250 (ALH)

German Research Foundation grant Li 690/14-1 (WAL)

IZKF Münster Li1/012/24 (WAL)

National Institutes of Health P41 GM103622(TCI), P30 GM138395 (TCI)

